# Mapping subcellular H_2_O_2_ dynamics reveals tissue specific redox patterns in *Drosophila*

**DOI:** 10.64898/2026.08.21.745691

**Authors:** Lucie A. G. van Leeuwen, Silvia Aldaz Casanova, Oisharja Rahman, Maura C. Hooiveld, Desiree M. C. Smakman, Jan Paul Lambooij, Tobias B. Dansen, Aniek Janssen, Hayley J. Sharpe

## Abstract

Redox signalling regulates development, tissue homeostasis, and organismal health. Hydrogen peroxide (H_2_O_2_) is a major signalling form of reactive oxygen species (ROS) that modulates protein activity through oxidation of redox-sensitive cysteines that is reversed by cellular reducing systems. Since H_2_O_2_ production, scavenging and reduction are spatially restricted, signalling specificity is strongly influenced by subcellular localisation. However, subcellular H_2_O_2_ dynamics in animal tissues remain poorly understood. To address this, we generated and validated *Drosophila melanogaster* lines expressing the ultrasensitive, ultrafast ratiometric H_2_O_2_ biosensor HyPer7 targeted to mitochondria, nucleus, cytosol, or plasma membrane. With its highly conserved metabolic and signalling pathways, tractable lifespan, and powerful genetic toolkit, *Drosophila* is an ideal model for studying redox biology. These new “FlyPer” lines enable tissue-specific HyPer7 expression and high-resolution measurement of subcellular, *in vivo* H_2_O_2_ dynamics throughout the lifespan. Using FlyPer, we detected compartment-specific H_2_O_2_ dynamics during oxidative stress, ageing, wing disc development and embryogenesis, uncovering unexpected patterns of spatially and temporally regulated oxidation throughout the organism. Together, these findings establish FlyPer as a valuable toolkit for *in vivo* redox biology and suggest that compartmentalised redox dynamics are a fundamental yet still poorly understood layer of developmental programming.

**Highlights:**

- FlyPer reports *in vivo* H_2_O_2_ dynamics across tissues and subcellular compartments from embryo to adult
- Redox states are differentially regulated across subcellular compartments during embryonic development
- Specific embryonic cell types display distinct developmental subcellular H_2_O_2_ dynamics
- Redox gradients mirror anatomical features and morphogen patterns

## Introduction

Hydrogen peroxide (H_2_O_2_) is an important signalling molecule that can regulate protein function through reversible oxidation of cysteine residues. Such oxidative modifications can influence numerous protein properties, including subcellular localisation, interactions with binding partners, enzymatic activity, and protein stability. This form of cysteine-oxidation dependent signal transduction, termed redox signalling, plays an essential role in many cellular processes, ranging from cell growth and metabolic signalling to immune cell activation and epigenetic regulation^1–3^. While other types of reactive oxygen species (ROS), such as the superoxide anion radical (O_2_^•−^), can also mediate redox signalling, H_2_O_2_ is considered particularly physiologically relevant due to its ability to cross membranes, high stability, low reactivity, and selectivity for cysteine thiols^4^.

Excessive H_2_O_2_ can lead to oxidative stress, which can cause oxidative damage to DNA, lipids and proteins^4^. Consequently, the production and removal of H_2_O_2_ must be tightly regulated to maintain redox homeostasis and ensure appropriate redox signalling^5,6^. H_2_O_2_ is directly produced by several oxidases, dehydrogenases and through dismutation of O_2_^•−^ by superoxide dismutases across various subcellular compartments, including the mitochondria, plasma membrane, cytosol, endoplasmic reticulum, peroxisomes and the nucleus^4,7,8^. Scavenging is predominantly carried out by the enzyme catalase, peroxiredoxins, and glutathione peroxidases, which show compartment-specific distribution^9^. The high efficiency of these scavenging systems typically restricts H_2_O_2_ actions to close vicinity of its production site^10–12^, thereby contributing to the specificity required to successfully fulfil its dynamic redox signalling functions. Another important determinant of redox signalling is the cellular reduction machinery, including the thioredoxin (Trx) and glutathione systems, which restores oxidised cysteine residues to their reduced thiol state and is itself subject to compartment-specific regulation and dependent upon NADPH supply^13–16^. Each location therefore hosts its own redox environment, with specific cross-talk between H_2_O_2_ producing enzymes, scavenging systems, reductants, and distinctive redox reactive protein networks^8^. Considering this, it is not difficult to envision that, for instance, H_2_O_2_ oscillations at the plasma membrane have different regulatory functions compared to H_2_O_2_ fluxes in the nucleus or the mitochondria^17^. Indeed, studies demonstrated that the cellular consequences of H_2_O_2_ fluxes vary depending on subcellular production site, and on whether it was intracellularly generated or exogenously applied^11,18–20^. Therefore, to study the downstream effects of H_2_O_2_-dependent redox signalling, it is not only important to know the amplitude and duration of the H_2_O_2_ signal, but also where it is situated within the cell.

Various approaches exist to measure intracellular H_2_O_2_ levels and dynamics in cells^10,21–29^. However, detecting physiological H_2_O_2_ signalling at the subcellular scale requires high-precision tools with sufficient spatial and temporal resolution. For this purpose, genetically encoded biosensors have been developed. Among these biosensors, HyPer7 is considered an ultrafast, ultrasensitive ratiometric probe that has been successfully employed to report on subcellular H_2_O_2_ dynamics both *in vitro* and in multiple model systems, such as yeast, worms, zebrafish, mice and rats^10,30,20,31–35^. It is one of the latest in a series of HyPer probes, and consists of a circularly permuted yellow fluorescent protein (YFP) integrated into a H_2_O_2_-sensitive OxyR domain from *Neisseria meningitidis*^10^. Unlike previous HyPer probes, it is pH stable and shows enhanced brightness. Following oxidation by H_2_O_2_, two reactive cysteine residues within the OxyR domain form an intramolecular disulfide bond. This redox-dependent conformational change increases the YFP emission at 516 nm when excited at its 499 nm excitation maximum, while reducing emission when excited at its 400 nm excitation maximum. Previous studies in yeast (*Saccharomyces cerevisiae*) show that the intramolecular disulfide bond in HyPer7 is predominantly reduced by the Trx system^31^. This has the important implication that HyPer7 does not simply measure the presence of H_2_O_2_. Rather it reports the balance between H_2_O_2_-driven oxidation and thiol reducing capacity of the Trx/TrxR/NADPH system within a specific cellular compartment. Consequently, HyPer7 more appropriately reflects a measure of H_2_O_2_ homeostasis, or redox dynamics, than abundance alone.

The fruit fly *Drosophila melanogaster* is an ideal model organism to study subcellular H_2_O_2_ signalling, owing to its high conservation of metabolic pathways and tissue function. Moreover, exceptionally powerful genetic tools for spatiotemporal control of gene expression and a vast collection of publicly shared mutant lines, enable the study of H_2_O_2_ signalling in complex *in vivo* contexts. Current methods to measure compartmentalised H_2_O_2_ in fruit flies include MitoB, which is a mitochondrial-targeted, ratiometric mass spectrometry probe that can be used to quantify mitochondrial H_2_O_2_ at the bulk tissue level^36^. In addition, transgenic flies expressing the ratiometric biosensor roGFP2-Orp1 have enabled the monitoring of H_2_O_2_ dynamics in both larval and adult tissues at subcellular resolution, with targeting to the cytosol and mitochondria^37^. While these transgenic fly lines represented an important advance, H_2_O_2_ measurement is restricted to a limited number of cellular compartments. Since their first incorporation in flies over a decade ago, roGFP2 probes and genetically encoded biosensors in general have undergone significant improvements to increase their sensitivity, response kinetics and dynamic range, with HyPer7 emerging as an important innovation. Additionally, roGFP2-based biosensors and HyPer7 are primarily reduced by distinct cellular reducing systems—the glutathione and thioredoxin systems, respectively—and can therefore provide complementary information on cellular redox homeostasis^31^. Altogether, we anticipate that flies expressing subcellular-targeted HyPer7 biosensors will provide a valuable resource for the comprehensive study of redox dynamics *in vivo*.

In this study, we have generated and validated four fly lines that each target HyPer7 to a different subcellular compartment: the mitochondrial matrix, nucleus, cytosol and inner leaflet of the plasma membrane. These fly lines carry their HyPer7 construct under an upstream activating sequence (UAS), allowing spatiotemporal control over HyPer7 expression using the widely used GAL4/UAS system in *Drosophila*^38^. We show that these “FlyPer” lines can be successfully used to measure *in vivo* H_2_O_2_ dynamics across cellular subcompartments, various tissues and developmental stages. We identify increases in subcellular H_2_O_2_ in flies fed with redox cycling compounds, and in ageing flies, indicating its sensitivity in adult flies *in vivo*. Moreover, our analysis reveals subcompartment-specific oxidation patterns of HyPer7 across larval imaginal discs and during embryogenesis. Collectively, these distinct patterns highlight dynamic subcellular regulation of oxidation by H_2_O_2_ and suggest potential roles for compartmentalised redox signalling in a variety of developmental processes.

## Results

### Generation of FlyPer lines

To determine how physiological H_2_O_2_ dynamics vary between different subcellular structures across tissues, cell types and developmental stages, an *in vivo* approach is essential. We therefore generated fly lines with the HyPer7 biosensor genetically integrated into their genome. We selected four subcellular compartments for targeting HyPer7: the mitochondrial matrix, nucleus, cytosol and inner leaflet of the plasma membrane (Fig. 1A). Each of these compartments hosts their own redox environment, which is crucial in maintaining cellular and organismal health^8^. The mitochondrial matrix is of interest because it is central to metabolic regulation and redox buffering, houses the mitochondrial genome, and is a key site of H_2_O_2_ production through O_2_^•−^ generation by the electron transport chain. The nuclear redox environment is an important regulator of gene expression and genome stability. The plasma membrane is a major signalling platform for receptor-mediated H_2_O_2_ responses and production via NADPH oxidases, and the cytosol serves as a central interface to integrate many H_2_O_2_ signalling and redox relay processes. It is therefore important to characterise H_2_O_2_ dynamics within each compartment *in vivo*, providing a foundation for understanding their contributions to tissue homeostasis and organismal health.

**Figure 1.**
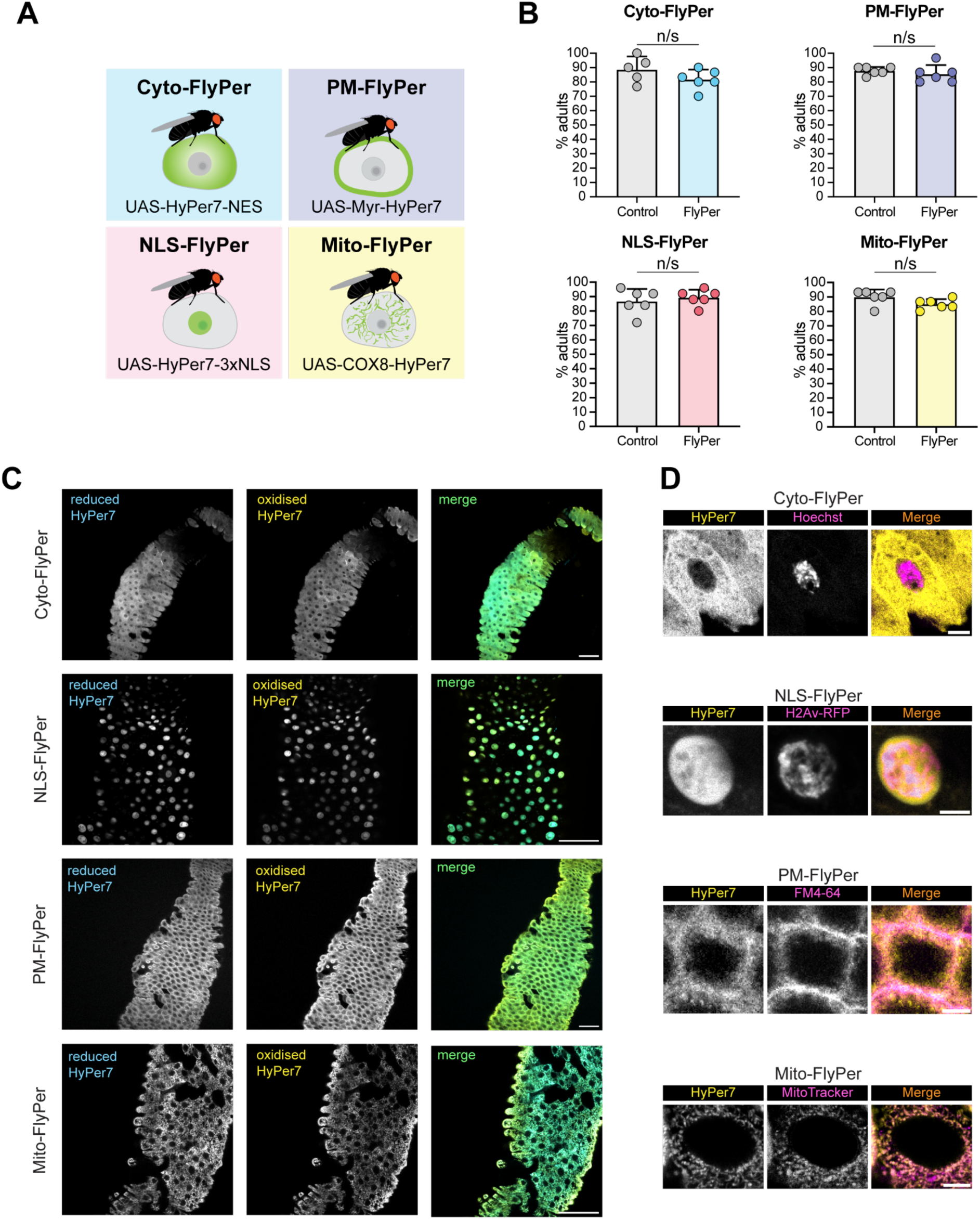
FlyPer lines are viable and show expected subcellular localisation. **(A)** Overview of the UAS-FlyPer lines generated in this study. For each subcellular compartment, an individual fly line was generated carrying the HyPer7 sequence fused to a subcellular localisation sequence, under the control of the UAS-promoter. Cytosol = Cyto-FlyPer; plasma membrane = PM-FlyPer; nucleus = NLS-FlyPer; mitochondrial matrix = Mito-FlyPer. **(B)** Bar graphs showing adult viability percentages of FlyPer lines ubiquitously expressing their HyPer7 construct using the *daughterless*-GAL4 driver (daGAL4). Heterozygous daGAL4 flies (crossed to *w^Dah^* wild type flies) were used as control genotype. The percentage of eclosed adults was determined from *n* = 5-6 vials each containing 25-30 embryos (150*-*180 embryos in total) for each genotype. Data are presented as mean ± SD, analysed by Welch’s t-test (n/s, *p* > 0.05). **(C)** Representative live confocal images of the adult midgut of female flies ubiquitously expressing FlyPer constructs (using daGAL4 driver) localising to distinct subcellular compartments, indicated on the left. Images show reduced HyPer7 (left column), oxidised HyPer7 (middle column) and the merged signal (right column). Scale bar = 50 µm. **(D)** Representative live confocal images showing subcompartmental localisation in the different FlyPer lines. Top shows cytosolic localisation of cyto-FlyPer (yellow) confirmed by nuclear exclusion (Hoechst, magenta) in midgut enterocyte. NLS-FlyPer (second from top, yellow) shows nuclear localisation confirmed by colocalisation with H2Av-RFP (magenta) in midgut enterocyte of flies co-expressing both transgenes (maximum projection). PM-FlyPer (second from bottom, yellow) shows localisation to plasma membrane confirmed by colocalisation with FM4-64 dye (magenta) in midgut enterocyte. Mito-FlyPer (bottom, yellow) shows localisation to mitochondria confirmed by MitoTracker Deep Red (magenta) in an adult female malpighian tubule stellate cell. Scale bar = 5 µm.

We first verified whether HyPer7 can be successfully expressed and employed to report on H_2_O_2_ fluctuations in fly cells in culture. We therefore expressed cytosolic HyPer7 (cyto-HyPer7) in cultured *Kc167* (*Kc*) cells by transient transfection (Fig. S1A). HyPer7 was readily detected in *Kc* cells and rapidly responded to exogenous H_2_O_2_ (Fig. S1B). We next tested whether cyto-HyPer7 also reacts to treatment with diamide. This thiol oxidising agent has previously been shown to increase HyPer7 oxidation^39^, as well as oxidation of the H_2_O_2_ sensor roGFP2-Orp1 in flies^37^, presumably by depleting reduced glutathione and oxidising Trx and other proteins tipping the balance towards more oxidation. Consistent with this, cyto-HyPer7 oxidation increased upon exposure to diamide (Fig. S1C).

We then proceeded to generate transgenic *Drosophila melanogaster* lines expressing HyPer7 constructs under a UAS promoter, termed “FlyPer” lines, enabling tissue- and cell type-specific expression through the widely used GAL4/UAS system^38^. We assessed viability in each FlyPer line ubiquitously expressing each subcellular HyPer7 construct (Fig. 1B). We observed no effects on developmental lethality in any of the lines (Fig. 1B), with adult viability ranging between ∼82%-90% for all genotypes. However, flies ubiquitously expressing mitochondrial HyPer7 (mito-HyPer7; mito-FlyPer) are developmentally delayed by approximately 1 day (Fig. S2A). This may indicate a modest sensitivity of mitochondrial redox homeostasis to constitutive HyPer7 expression. In contrast, flies expressing plasma membrane-targeted HyPer7 (PM-HyPer7; PM-FlyPer), nuclear HyPer7 (NLS-HyPer7; NLS-FlyPer) or cytosolic HyPer7 (cyto-FlyPer) exhibit development timing comparable to control flies (Fig. S2B-D).

Finally, using the ubiquitous drivers, daughterless-GAL4 (daGAL4) or tubulin-GAL4 (tubGAL4), we observed stable expression of all HyPer7 constructs across every *Drosophila* tissue tested. Live imaging of the adult midgut, one of the primary tissues examined in this study, revealed the distinct subcellular localisation patterns of each HyPer7 reporter (Fig. 1C). Co-localisation analysis with established markers for the four targeted compartments further confirmed that each HyPer7 construct was accurately and consistently localised to their intended subcellular location (Fig. 1D). Together, these data demonstrate that HyPer7 can detect H_2_O_2_ dynamics in fly cells. FlyPer lines ubiquitously expressing HyPer7 are furthermore viable and display expected subcellular localisation.

### FlyPer lines detect ex vivo H_2_O_2_

We next verified if HyPer7 sensors appropriately respond to elevated H_2_O_2_ in tissues. To do this, we exposed different dissected tissues of various FlyPer lines to a range of H_2_O_2_ concentrations. First, we rapidly dissected the adult midgut of female flies ubiquitously expressing cyto-HyPer7. Guts were then treated with or without 250 µM H_2_O_2_ in the dissection medium for 15 minutes, followed by immediate live confocal imaging (Fig. 2A). Cyto-HyPer7 oxidation significantly increased upon H_2_O_2_ treatment, indicating successful detection of exogenous H_2_O_2_ in a tissue *ex vivo* (Fig. 2B). We then investigated whether HyPer7 could detect intracellular accumulation of H_2_O_2_ following diamide treatment *ex vivo*, as previously observed in cultured *Kc* cells (Fig. S1C). Live imaging was performed on guts dissected from NLS-FlyPer flies, with analyses focussed on midgut enterocytes. These cells are readily identified by their highly polyploid state, which gives rise to enlarged nuclei^40,41^. Indeed, diamide induced a marked increase in NLS-HyPer7 oxidation (Fig. 2C, D). Next, we assessed responses in the mitochondria of stage 9-10 egg chambers (Fig. 2E) using live imaging. While treatment with 10 µM H_2_O_2_ increased oxidation over a 30 minute timeframe (Fig. 2F), 100 µM led to an initially robust response at 6 minutes, which subsequently decreased at 30 minutes (Fig. 2G), consistent with HyPer7 reduction. Collectively, these findings demonstrate that the FlyPer lines can reliably detect exogenous H_2_O_2_ across various compartments and *ex vivo* tissues, as well as reporting on cellular reductive capacity.

**Figure 2.**
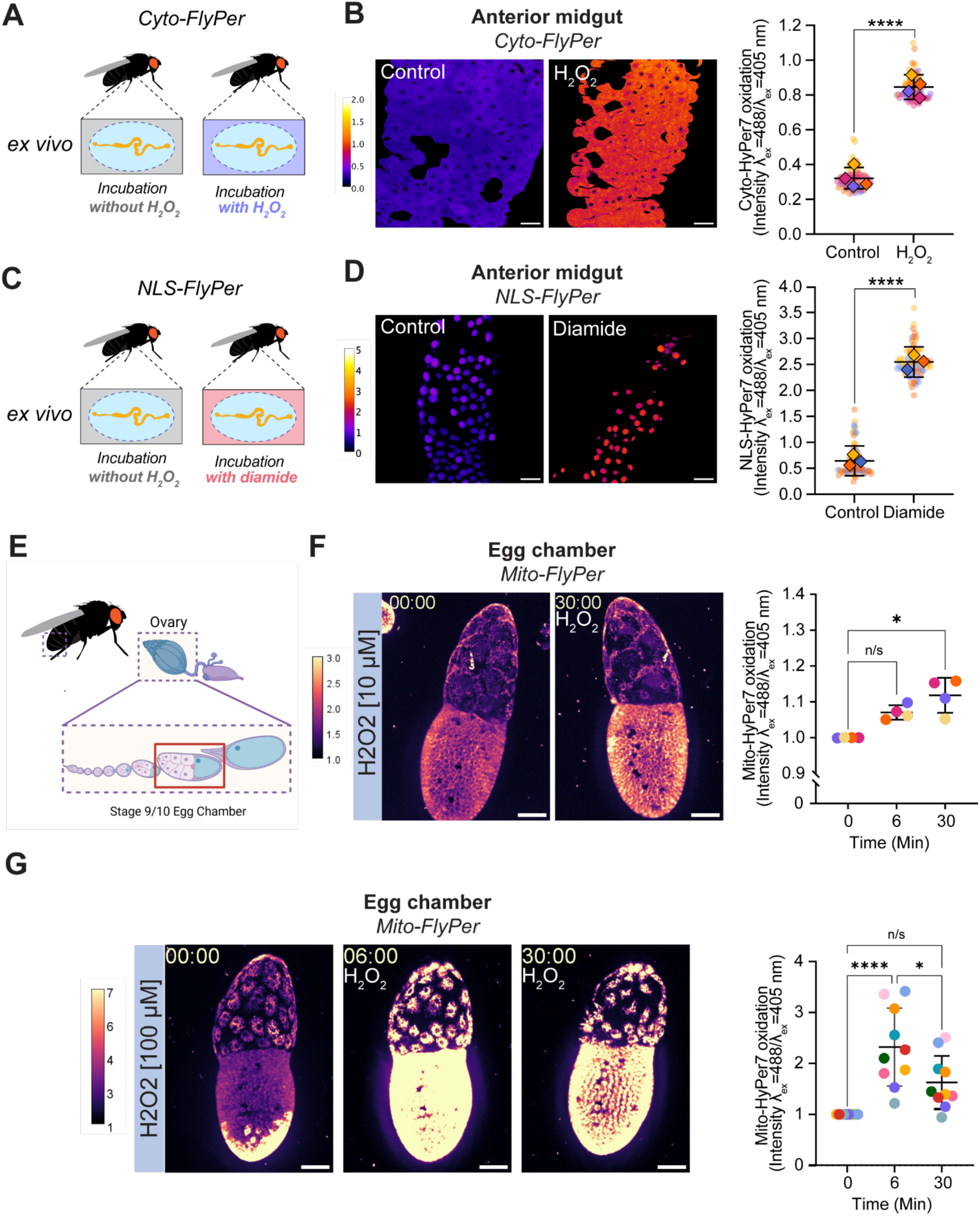
FlyPer lines detect *ex vivo* oxidation by H_2_O_2_ across tissues and subcellular compartments. **(A)** Schematic of dissected adult midguts of cyto-FlyPer flies (daGAL4>UAS-HyPer7-NES) treated with either medium (CCM3 medium) or exogenous H_2_O_2_ (250 µM in CCM3 medium) for 15 minutes before being live imaged by confocal microscopy. (**B**) (Left) Representative ratiometric images of a control gut (top) and H_2_O_2_-treated gut (bottom) dissected from cyto-FlyPer flies. (Right) Quantification of oxidised/reduced cyto-HyPer7 signal in adult midgut enterocytes without (Control) or with (H_2_O_2_) exposure to exogenous H_2_O_2_ *ex vivo*. Data are presented as means (diamond shapes) of ∼20 cells (dots) per gut, with the horizontal line indicating the overall mean ± SD. Data were analysed by Welch’s t-test (****, *p* < 0.0001). *n* = 4 guts per condition. **(C)** Schematic of dissected adult midguts of NLS-FlyPer flies (daGAL4>UAS-HyPer7-3xNLS) treated with either medium (CCM3 medium) or diamide (1 mM in CCM3 medium) for 20 minutes before being live imaged by confocal microscopy. **(D)** (Left) Representative ratiometric image of a control gut (top) and a diamide-treated gut (bottom) dissected from NLS-FlyPer flies. (Right) Quantification of oxidised/reduced NLS-HyPer7 signal in adult midgut enterocytes without (Control) or with (Diamide) *ex vivo* diamide treatment. Data are presented as means (diamond shapes) of ∼20 cells (dots) per gut, with the horizontal line indicating the overall mean ± SD. Data were analysed by Welch’s t-test (****, *p* < 0.0001). *n* = 3 guts per condition. Scale bar = 25 µm. **(E)** Schematic of fly ovary dissection and egg chamber stages, where stage 9/10 ovaries were selected (highlighted by red box). Schematic created using Biorender.com. **(F)** (Left) Representative ratiometric images of stage 10 egg chamber, at timepoint 0 and 30 minutes post addition of exogenous H_2_O_2_ (10 µM in Schneider’s Insect medium) *ex vivo* from mito-FlyPer flies (tubGAL4>UAS-COX8-HyPer7). Scale bar = 50 µm. (Right) Quantification of oxidised/reduced mito-HyPer7 signal in egg chambers pre- (Control) or post (H_2_O_2_) treatment *ex vivo*. Each data point represents one egg chamber, and the horizontal line indicates mean ± SD. Data were analysed by Friedman test (*, *p* < 0.05; n/s, *p* > 0.05). *n* = 4 egg chambers. **(G)** (Left) Representative ratiometric images of stage 9-10 egg chambers, at timepoints 0, 6 and 30 mins post addition of exogenous H_2_O_2_ (100 µM in Schneider’s Insect medium) *ex vivo* from mito-FlyPer flies. Scale bar = 50 µm. (Right) Quantification of oxidised/reduced mito-HyPer7 signal in egg chambers pre- (Control) or post (H_2_O_2_) treatment *ex vivo* at each time point (0, 6 and 30 minutes). Each data point is coloured to correspond to the same individual egg chamber and the horizontal line indicates mean ± SD. Data were analysed by Friedman test for repeated measures (****, *p* < 0.0001; *, *p* < 0.05; n/s, *p* > 0.05). *n* = 10 egg chambers.

### Mito-FlyPer line detects in vivo H_2_O_2_ fluctuations following paraquat-induced stress

Having established that we can detect H_2_O_2_ fluctuations *ex vivo*, we then explored whether FlyPer reports on *in vivo* dynamics. The redox cycler paraquat is frequently used to induce mitochondrial superoxide production, including in flies^42^. This superoxide is subsequently converted to H_2_O_2_ by manganese superoxide dismutase (mnSOD) in the mitochondrial matrix^43–45^. Using the ratiometric mass spectrometry probe mitoB, it was previously demonstrated that paraquat feeding results in increased H_2_O_2_ levels in fly mitochondria^46^.

To test if dietary treatment with 20 mM paraquat leads to increased mito-HyPer7 oxidation, we fed 7-day old mito-FlyPer flies either their regular diet or paraquat-supplemented food for 24 h. Since the gut is among the first tissues to be exposed to paraquat upon food ingestion, we then dissected and immediately imaged fly midguts by live confocal microscopy. We observed a substantial increase in oxidised mito-HyPer7 in the midgut of paraquat-fed flies in comparison to their control cohort (Fig. 3A, B). To determine whether mitochondrial H_2_O_2_ dynamics can be monitored in tissues exposed to paraquat through systemic circulation, we performed live imaging of the adult fat body of control and paraquat-fed flies. The *Drosophila* adult fat body is equivalent to vertebrate adipose tissue and liver, and is an important tissue in regulating energy metabolism, immunity and reproduction^47^. Although the fat body plays an established role in the oxidative stress response^48–50^, the effects of paraquat on mitochondrial H_2_O_2_ dynamics have not previously been examined directly in this tissue. Mito-HyPer7 oxidation increased significantly in the fat body following paraquat treatment (Fig. 3C, D), demonstrating an expanded response to paraquat, but also that the reporter is sufficiently sensitive to detect compartment-specific oxidative responses to *in vivo* challenges.

**Figure 3.**
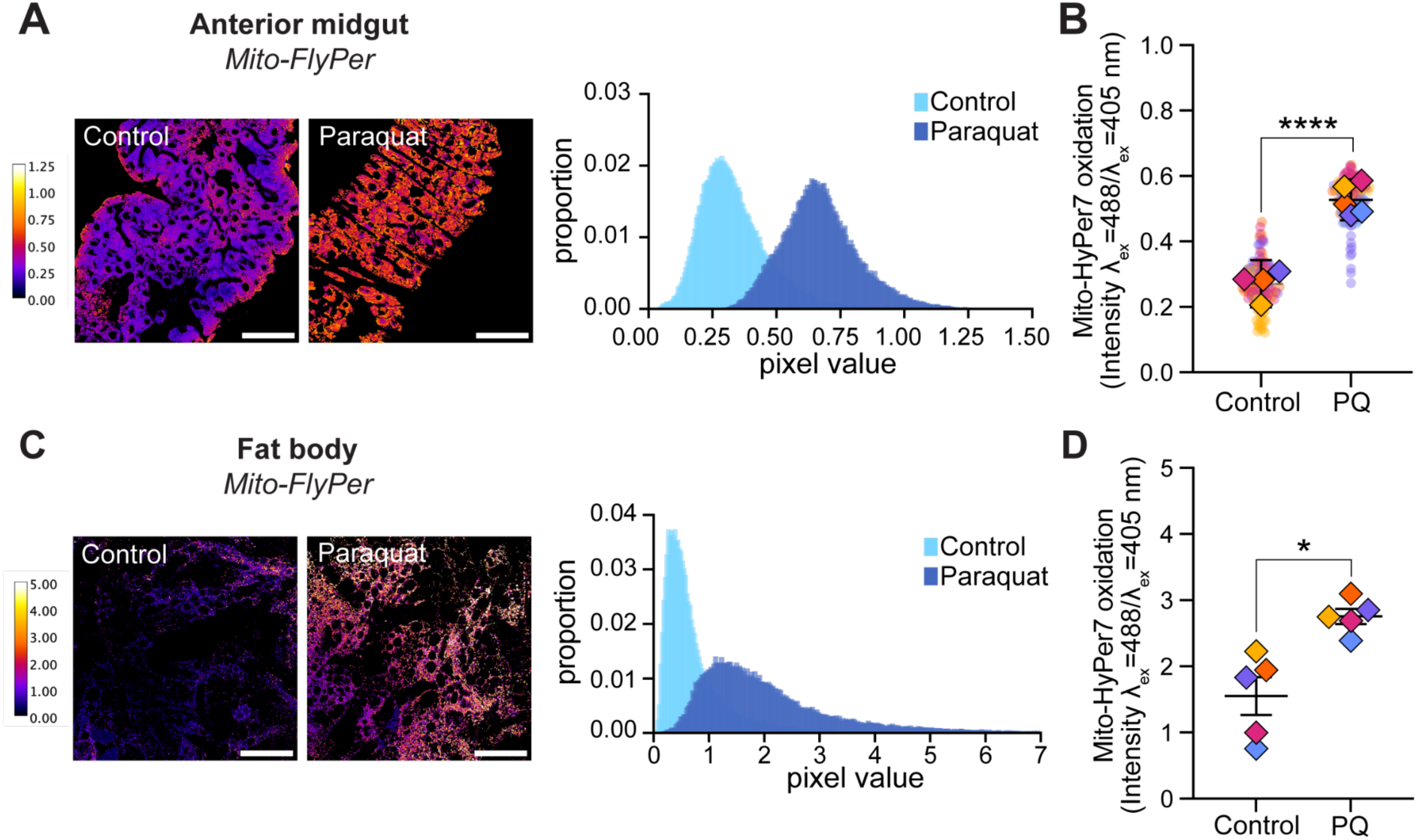
Mito-FlyPer detects increased mitochondrial H_2_O_2_ in the gut and fat body of paraquat- treated flies. **(A)** Representative ratiometric images of anterior midguts of 7-day old female mito-FlyPer flies (daGAL4>UAS-COX8-HyPer7) kept on their standard diet (Control) or a paraquat-supplemented diet (Paraquat) for 24 h. The histogram shows the distribution of pixel values in the depicted ratiometric images. Scale bar = 50 µm **(B)** Quantification of anterior midguts of female mito-FlyPer flies (daGAL4>UAS-COX8-HyPer7) kept on their standard diet (Control) or a paraquat-supplemented diet (PQ) for 24 h. Data are presented as means (diamond shapes) of ∼30 cells (dots) per gut, with the horizontal line indicating the overall mean ± SD. Data were analysed by Welch’s t-test (****, *p* < 0.0001). *n* = 4-5 guts per condition. **(C)** Representative ratiometric images of fat body tissue of 7-day old mito-FlyPer flies (daGAL4>UAS-COX8-HyPer7) kept on their standard diet (Control) or a paraquat-supplemented diet (Paraquat) for 24 h. The histogram shows the distribution of pixel values in the depicted ratiometric images. Scale bar = 100 µm. **(D)** Quantification of fat bodies of 7-day old mito-FlyPer expressing flies (daGAL4>UAS-COX8-HyPer7) kept on their standard diet (Control) or a paraquat-supplemented diet (PQ) for 24 h. Data are presented as means (diamond shapes) of 2 regions per fat body, with the horizontal line indicating the overall mean ± SD. Data were analysed by Welch’s t-test (*, *p* < 0.05), *n* = 5 fat bodies per condition.

### Cyto-FlyPer detects physiological H_2_O_2_ dynamics in ageing

Paraquat-induced redox stress represents an acute and severe challenge that drives elevated mitochondrial H_2_O_2_. We therefore asked whether the FlyPer lines could also report physiologically relevant H_2_O_2_ fluctuations under more natural conditions. Ageing provides such a context, as cytosolic H_2_O_2_ levels have been shown to increase in the female *Drosophila* midgut, as reported using the genetically encoded H_2_O_2_ probe roGFP-Orp1^37^. We tested whether our novel cyto-FlyPer line could be used to replicate this finding. As an initial control, we verified that expression of cyto-HyPer7 did not adversely affect healthy ageing. Female flies ubiquitously expressing cyto-HyPer7 under the control of the daGAL4 driver were, if anything, slightly longer lived compared to control flies (Fig. S3), indicating HyPer7 expression does not impair lifespan. We then compared the oxidation state of cyto-HyPer7 in the anterior midgut of young (7-day old) and aged (56-day old) cyto-FlyPer females. Consistent with previous observations, cyto-HyPer7 was modestly, but reproducibly more oxidised aged flies (Fig. 4A, B). Together, these results validate the ability of the FlyPer lines to detect physiologically relevant, endogenous changes in H_2_O_2_ levels during ageing and highlight their utility for investigating this in other, unexplored biological and subcellular contexts.

**Figure 4.**
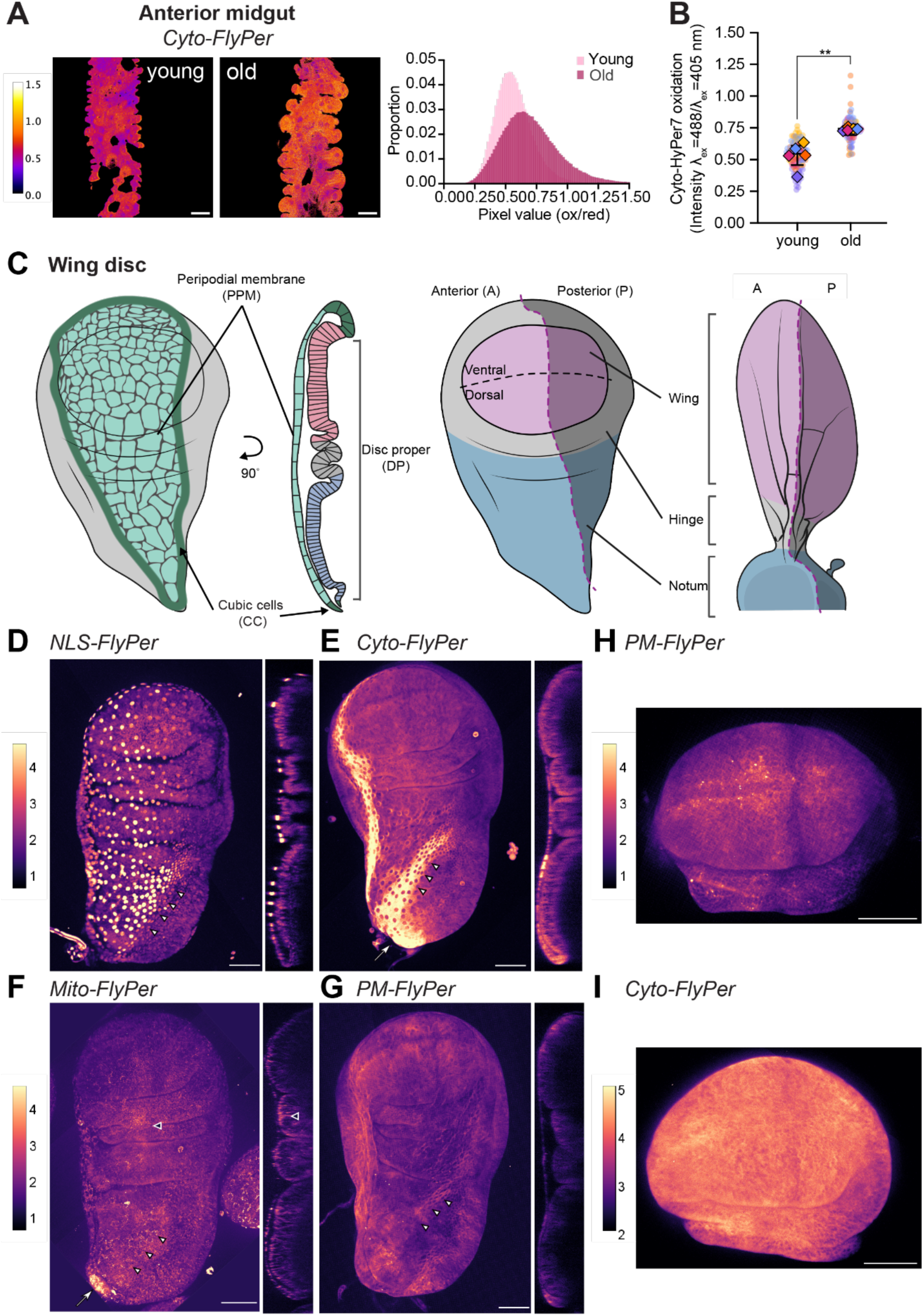
FlyPer lines reveal physiological H_2_O_2_ dynamics across subcellular compartments in the larval wing disc. **(A)** Representative ratiometric images of cyto-FlyPer (daGAL4>UAS-HyPer7-NES) in the anterior midgut of young (day 7) and old (day 56) female flies. The histogram shows the distribution of pixel values in the depicted ratiometric image. Scale bar = 25 µm. **(B)** Quantification of young (day 7) and old (day 56) female cyto-FlyPer expressing flies (daGAL4>UAS-HyPer7-NES) in the anterior midgut. Data are presented as means (diamond shapes) of ∼15-20 cells (dots) per gut, with the horizontal line indicating the overall mean ± SD. Data were analysed by Welch’s t-test (**, *p* < 0.01), *n* = 5 midguts per age group. **(C)** Schematic of third instar larval wing disc. Left: en face and orthogonal views of the peripodial membrane (PPM) and cuboidal cells (CC; green) that connect to the disc proper (DP). Right: Disc proper and its relationship to the adult wing. The dorsal-ventral and anterior-posterior boundaries act as organising centres that control tissue growth, patterning and organisation. **(D-G)** Representative ratiometric en face (left) and orthogonal cross section (right) images of the third instar imaginal wing disc of (D) flies expressing NLS-FlyPer (tubGAL4>UAS-HyPer7-3xNLS), (E) flies expressing cyto-FlyPer (tubGAL4>UAS-HyPer7-NES), (F) flies expressing mito-FlyPer (tubGAL4>UAS-COX8-HyPer7) and (G) flies expressing PM-FlyPer (tubGAL4>UAS-Myr-HyPer7). Scale bar = 50 µm. Arrow indicates increased HyPer7 oxidation in the stalk that is severed to isolate discs for imaging. White triangles indicate the CC. Black triangles indicate increased HyPer7 oxidation in the hinge fold region. **(H-I)** Representative ratiometric images of the wing pouch from fly larvae expressing (H) PM-FlyPer (nubGAL4>UAS-Myr-HyPer7) or (I) cyto-FlyPer (nubGAL4>UAS-HyPer7-NES) using a DP-specific driver. Scale bar = 50 µm.

### FlyPer lines reveal physiological subcellular H_2_O_2_ dynamics in the larval wing disc

To this end, we next explored subcellular H_2_O_2_ dynamics using all four FlyPer lines in the third-instar larval wing disc, which is a well-established model for studying growth and patterning^51,52^. Wing discs are epithelial sacs composed of two layers: a squamous epithelial layer called the peripodial membrane (PPM), and a pseudostratified columnar epithelial layer that forms the disc proper (DP) and eventually contributes to the majority of the adult cuticular structures of the wing (Fig. 4C)^53^. Cells in the PPM form a monolayer with large, flat nuclei, while the cells in the DP are columnar with long, stretched, intercalated nuclei. These PPM and DP layers are connected by a transitional region in which cells gradually change shape from columnar to squamous; these cells are referred to as cubic cells (CC; Fig. 4C).

NLS-FlyPer wing discs showed distinct patterns of nuclear oxidation; with higher levels in the PPM and CC (Fig. 4D) and lower and homogeneous levels in the DP (Fig. 4D). In contrast, wing discs from cyto-FlyPer flies showed higher levels of oxidation in the CC compared to the PPM in general (Fig. 4E). High levels of cyto-HyPer7 oxidation were also consistently observed at the ‘stalk’, the region that has to be severed to remove the disc from the larva, and could therefore reflect tissue injury (Fig. 4E, arrow). We note similar elevated levels of oxidation in this region for mito-FlyPer wing discs (Fig. 4F, arrow), but not in NLS-FlyPer or PM-FlyPer lines (Fig. 4D, G). In mito-FlyPer wing discs, modest but consistent higher levels of oxidation are observed around the hinge folds compared to the rest of the DP (Fig. 4F, black triangles). For PM-FlyPer, the DP shows higher apical oxidation as well as oxidation in the CC (Fig. 4G). The most striking feature for PM-FlyPer was revealed by imaging the wing pouch using nubbin-GAL4 to drive expression only in the DP (Fig. 4C, right). Here PM-HyPer7 is reduced specifically in the dorso-ventral and anterio-posterior borders, with a gradient that is more oxidised at distal (central) regions and more reduced at proximal regions (peripheral) (Fig. 4H). In contrast, cyto-FlyPer wing discs show some oxidation along the dorso-ventral but not the antero-posterior border (Fig. 4I).

The observed oxidation and reduction patterns in our various FlyPer lines suggest that H_2_O_2_ homeostasis in the wing disc is not uniform, but instead reveals interesting subcellular patterns aligning with specific anatomical features. Moreover, these intrinsic differences highlight the sensitivity of FlyPer lines in revealing physiologically relevant redox dynamics.

### In vivo imaging reveals subcellular H_2_O_2_ dynamics in FlyPer embryos

So far, we have used our FlyPer lines to study subcellular H_2_O_2_ dynamics in dissected tissues. *Drosophila*, however, also provides powerful opportunities for live, *in vivo* imaging. Embryos are immobile, optically accessible, genetically tractable and develop rapidly, making them an excellent system for long-term imaging studies. We therefore characterised FlyPer reporters during embryonic development (Fig. 5A). Strikingly, the different lines showed distinct spatiotemporal patterns of oxidation. PM-FlyPer exhibited the most dynamic redox behaviour during embryogenesis (Fig. 5B, movie S1). From the beginning, we observed maternally deposited yolk granules with high levels of PM-HyPer7 oxidation, persisting until dorsal closure and preceding embryonic transcription driven by tub-GAL4 (Fig. 5C; left). Notably, oxidised yolk granules were not detected in other FlyPer lines (Fig. 5D-F). Together this indicates the maternal contribution of PM-HyPer7 and reflects compartment-specific, specifically high plasma membrane content, rather than autofluorescence. During stages 14 to 15, elevated membrane oxidation was apparent at the leading-edge during dorsal closure and during the convergence and sealing of the epidermal sheets at the dorsal midline (Fig. 5C). Membrane oxidation also increased progressively throughout development, particularly within epidermal cells. At later stages of embryogenesis, widespread expression made it difficult to distinguish individual cell types, however, the gut and the posterior spiracles (PS) were identifiable and showed high levels of oxidation (Fig. 5B).

**Figure 5.**
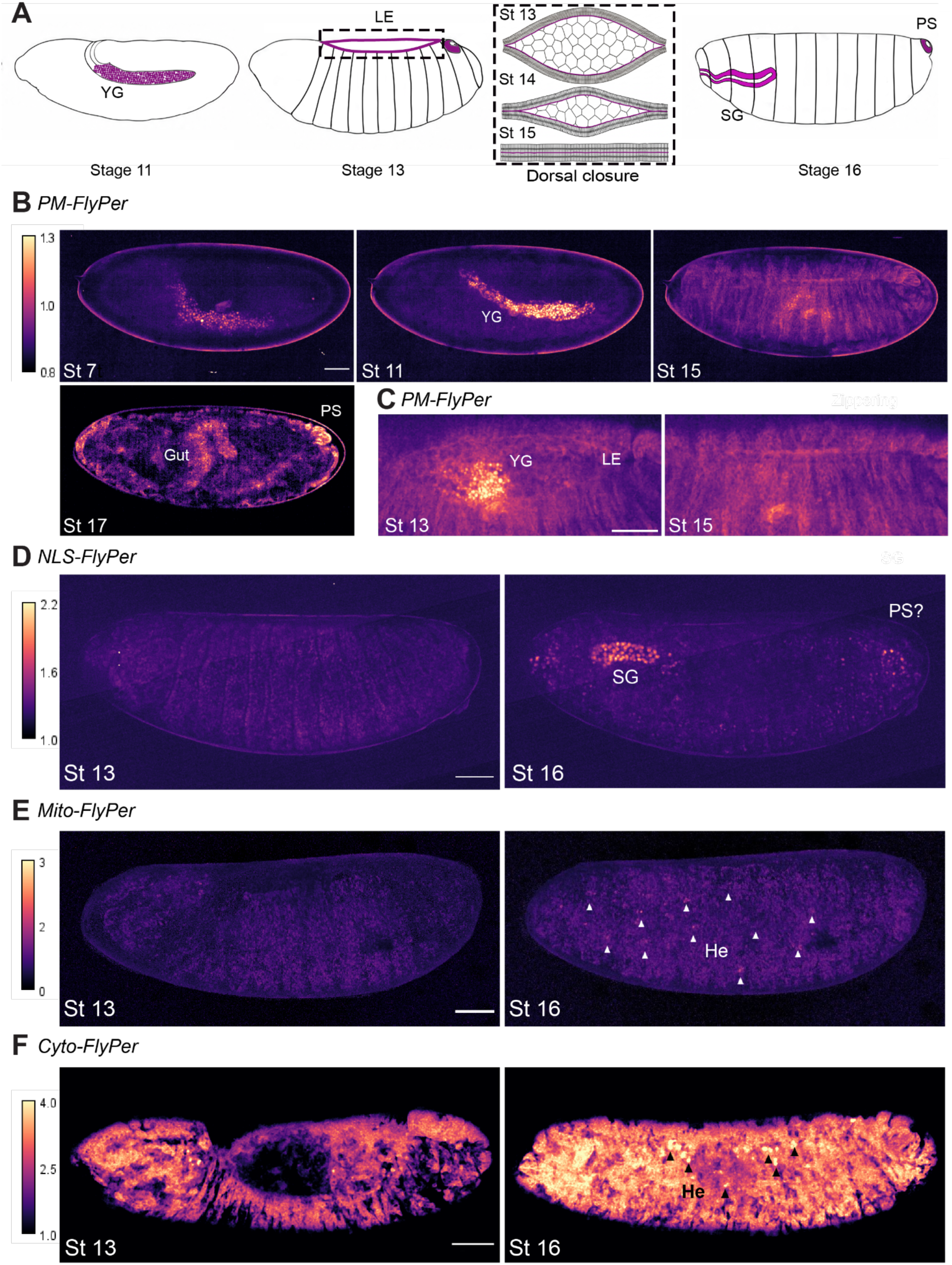
FlyPer lines reveal subcompartmental H_2_O_2_ dynamics during *Drosophila* embryogenesis. (**A**) Schematic representation of embryonic stages. From left to right: Embryo at stage 11 with the germ band fully extended. Yolk granules (YG) in purple. Stage 13 embryo during dorsal closure. Leading edge (LE) in purple. Representation of dorsal closure during stages 13-15 where columnar cells close over amnioserosa squamous cells. Leading edge in purple. Stage 16 embryo with salivary glands (SG) and posterior spiracles (PS) in purple. (**B**) Representative ratiometric images from PM-FlyPer (tubGAL4>UAS-Myr-HyPer7) expressing embryos (stages 7-17). Each image is an average projection of a Z stack. Scale bar = 50 μm. (**C**) Representative ratiometric images from PM-FlyPer expressing embryos focusing on the dorsal region at stage 13 (left) and stage 15 (right). Scale bar = 20 μm. (**D-F**) Representative ratiometric images from embryos of flies expressing NLS-FlyPer (D; tubGAL4>UAS-HyPer7-3xNLS), flies expressing mito-FlyPer (E; tubGAL4>UAS-COX8-HyPer7) and flies expressing cyto-FlyPer (F; tubGAL4>UAS-HyPer7-NES) at stage 13 (left) and 16 (right). White (E) and black (F) arrow heads indicate putative hemocytes (He). Each image is an average projection of a Z stack. Scale bar = 50 μm. Images are representative of *n* = 5 embryos per genotype.

In contrast, NLS-FlyPer embryos show very little evidence of dynamic oxidation during early embryogenesis. By stage 14, however, the nuclei of the salivary glands become strikingly oxidised (Movie S2). At later stages (stage 16), additional patches of cells in the head and tail, potentially corresponding to the posterior spiracles (PS), also displayed pronounced nuclear oxidation (Fig. 5D, right). Further studies will be required to identify these cells and to understand the mechanisms and functional significance of these localised nuclear oxidation events. Mito-FlyPer embryos exhibited homogeneous levels of oxidation across tissues throughout development (Fig. 5E, Movie S3). Nevertheless, a small population of motile cells beneath the epidermis showed elevated mito-HyPer7 oxidation, potentially corresponding to infiltrating hemocytes. In contrast, cyto-FlyPer embryos showed a progressive increase in cytoplasmic H_2_O_2_-dependent oxidation during development (Fig. 5F, movie S4). We also observed motile cells with elevated oxidation, reminiscent of those detected in mito-FlyPer embryos and likely representing a subset of hemocytes with distinct redox states. Collectively, these findings uncover previously unappreciated compartment, cell-type and tissue-specific patterns of H_2_O_2_-dependent oxidation during embryogenesis.

## Discussion

In this study, we established the first HyPer7-based Drosophila toolkit for monitoring H_2_O_2_ dynamics in vivo. We generated transgenic reporter lines targeting HyPer7 to the nucleus, cytosol, mitochondria, and plasma membrane, revealing striking compartment-specific patterns of oxidation across multiple tissues and developmental stages. By combining the sensitivity of HyPer7 with the genetic tractability of Drosophila, FlyPer provides a powerful platform for investigating H_2_O_2_-dependent signalling in development, ageing, and disease. More broadly, this work not only delivers a toolkit for visualising previously inaccessible redox signalling events *in vivo,* but also reveals that redox dynamics are an integral component of developmental programmes. The unexpected diversity, specificity, and dynamic nature of spatiotemporal oxidation patterns uncovered by FlyPer suggest that we are only beginning to appreciate the scope of redox regulation in living systems. We anticipate that these tools will accelerate the discovery of fundamental mechanisms and principles linking redox dynamics to cellular behaviour, tissue morphogenesis, homeostasis.

To demonstrate the utility of FlyPer, we monitored subcellular H_2_O_2_ dynamics across multiple physiological contexts. Using mito-FlyPer, we observed increased oxidation in the midgut and the fat body following paraquat exposure, extending previous reports of elevated ROS^54–56^. Previous studies relied on general ROS indicators, such as dihydroethidium (DHE) or CellROX, which lack subcellular resolution and suffer from issues of specificity^29^. While paraquat is known to induce mitochondrial oxidative stress, it can also affect H_2_O_2_ homeostasis in other compartments^45,57,58^. FlyPer will therefore be valuable for systematically addressing cell type- and compartment-specific responses to paraquat and other stressors.

We further validated FlyPer lines as reporters of physiological H_2_O_2_ fluctuations by imaging aged female midguts. Consistent with previous findings^37^, we observed an increase in cyto-HyPer7 oxidation with age, illustrating the suitability of these reporters for studying redox alterations during ageing. Our analyses of imaginal discs and embryos revealed diverse redox microenvironments during development. Although redox processes are known to influence embryogenesis and tissue patterning, the compartmentalised nature of redox signalling in this context is relatively unexplored^3,59,60^. Our observations are based primarily on morphological landmarks, thus co-localisation with additional markers or use of cell-type specific drivers will be required for further detailed characterisation. Nevertheless, several notable patterns of distinct redox states were apparent.

Mitochondria are a major source of intracellular H_2_O_2_^61,62^ and possess robust antioxidant buffering mechanisms, including peroxiredoxins and reductases^8,63^. In egg chambers, a high concentration of exogenous H_2_O_2_ led to rapid oxidation of mito-HyPer7 followed by reduction potentially through activation of adaptive antioxidant responses, for example via Nrf2 activity^7,64^. Under basal conditions, mitochondrial oxidation was relatively uniform across larval wing discs and embryonic tissues, consistent with tight redox control. However, increased mito-HyPer7 oxidation was detected in dispersed cells of stage 16 embryos, which are likely to represent a subset of activated hemocytes that use ROS analogously to macrophages and neutrophils^65,66^. A similar signal was observed in cyto-FlyPer embryos, but not from the other FlyPer lines. Further studies with specific markers are needed to better identify these cells and the function of hemocyte oxidation in development. A second pattern was increased oxidation in the wing disc stalk region in both cytosolic and mitochondrial reporter lines. Since this structure must be severed during dissection, this signal may reflect wound-induced H_2_O_2_ signalling. H_2_O_2_ and ROS are known mediators of responses to tissue damage^67^ regeneration^68^ and immune cell recruitment^69^ raising the possibility that FlyPer could be used to study the compartmentalisation of H_2_O_2_ kinetics and subcellular gradients during wound healing in more detail. Consistently with this, PM-FlyPer revealed elevated oxidation at the plasma membrane of leading edge cells during embryonic dorsal closure, which shares many features with wound healing^70,71^. Previous studies in *Drosophila* and zebrafish have highlighted an important role for H_2_O_2_ production by NADPH oxidase DUOX at the plasma membrane during tissue injury and repair^72,73^. Our observations suggest that similar mechanisms occur during developmental morphogenesis.

PM-FlyPer also revealed striking patterns in wing discs, indicating reduction at the dorso-ventral and anterior-posterior borders compared to the rest of the disc and adjacent gradients of oxidation. These regions are major signalling centers, controlling both growth and differentiation^74^. Together with the dynamic oxidation patterns observed during dorsal closure, our findings indicate that PM-FlyPer will be particularly valuable for identifying localized H_2_O_2_ fluxes during development and regeneration. NLS-FlyPer revealed elevated nuclear oxidation in embryonic salivary glands during embryogenesis and in the peripodial membrane nuclei of imaginal wing discs. To our knowledge, such patterns have not been described previously. Nuclear H_2_O_2_ can be produced by enzymes such as NADPH oxidase 4 and lysine-specific histone demethylase 1^75,76^, both conserved in *Drosophila,* and FlyPer now provides a means to investigate how such enzymes contribute to nuclear redox signalling during development

Although FlyPer is a powerful tool for monitoring compartment-specific H_2_O_2_ dynamics, several considerations should be noted. Despite generally normal viability, HyPer7 expression may influence physiology. For example, mito-FlyPer flies exhibited a one-day developmental delay, while cyto-FlyPer females showed a small lifespan extension. These effects may result from scavenging of endogenous H_2_O_2_ or increased demand on cellular reducing pathways. A previous study in yeast found that expressing HyPer7 had limited impact on the total cellular H_2_O_2_ scavenging capacity^31^, nevertheless, such issues can be mitigated using tissue-specific or inducible (*e.g.* GeneSwitch^77^ auxin-inducible lines^78^or temperature-sensitive drivers^79^) driver lines.

As recommended for previous H_2_O_2_-probes^80^, ratiometric measurements should be interpreted alongside individual (405 nm and 488 nm) channels to confirm reciprocal changes in HyPer7 emission and exclude imaging artefacts. HyPer7 is considerably less sensitive to intracellular pH fluctuations compared to previous HyPer variants^10,30,31^, reducing a major confounder in redox imaging. Likewise, differences in expression levels are generally not problematic because they affect both channels equally, although regions with very low reporter expression may need to be excluded from analyses. A further consideration is phototoxicity, since confocal imaging can lead to ROS generation^81^. During embryo imaging, mito and cyto-FlyPer reporters were more sensitive to imaging conditions than NLS- and PM-FlyPer. Under spinning disk imaging conditions, embryos expressing cyto- or mito-HyPer7 did not survive 16 h imaging (*n* = 5 embryos for cyto-FlyPer and *n* = 4 embryos for mito-FlyPer), whereas they remained viable under the confocal imaging conditions described in the methods. These observations highlight the importance of minimising laser power and exposure times during long-term live imaging.

The observed patterns of HyPer7 oxidation reflect a dynamic combination of local H_2_O_2_ concentrations and the activity of Trx/TrxR/NADPH-dependent thiol-reducing machinery. Consequently, HyPer7 oxidation patterns of interest should be viewed as integrated readouts of local redox homeostasis. The extensive *Drosophila* toolkit provides numerous possibilities to manipulate H_2_O_2_-producing enzymes, antioxidant systems and signalling pathways. These studies can be complemented by pharmacological treatments, redox proteomics, targeted approaches to assess cysteine oxidation in candidate proteins, and redox-inert knock-in mutants to determine the molecular consequences of H_2_O_2_-based signalling^82–87^.

## Acknowledgements

We thank all Janssen, Sharpe and Lens lab members for input on experimental design during laboratory meetings, as well as Ian McGough (Babraham Institute) for advice on *Drosophila* construct design, Simon Walker, Anneliese Jarman and Isabel San Martin Molina from Babraham Institute Imaging facility and Tiffany Lai for schematics. We furthermore gratefully acknowledge the Bloomington *Drosophila* Stock Center (NIH P40OD018537). This work was supported by an Engineering and Physical Sciences Research Council (ESPRC) Frontier guarantee grant (EP/Z000114/1), a Biotechnology and Biological Sciences Research Council (BBSRC) institutional programme grant (BBS/E/B/000C0433), core capability grant (BB/CCG2210/1) and a Netherlands Research Council (NWO) ENW-M1 grant (OCENW.M.22.347). OR is supported by a studentship funded by the Lister Institute of Preventive Medicine. HJS is an EMBO (European Molecular Biology Organization) Young Investigator and Lister Institute Prize Fellow. LAGvL is supported by a Netherlands Research Council (NWO) XS grant (OCENW.XS22.4.138).

## Author contributions

Conceptualisation: LAGvL, OR, TBD, AJ, and HJS.

Investigation: LAGvL, SAC, OR, MCH, DMCS, and JPL.

Formal analysis: LAGvL, SAC, and OR.

Resources, TBD, AJ and HJS.

Writing – original draft: LAGvL, SAC, and OR.

Writing – review and editing: LAGvL, TBD, AJ, and HJS.

Supervision: LAGvL, SAC, TBD, AJ, and HJS.

Funding acquisition: LAGvL, AJ, and HJS.

Project administration: AJ and HJS.

## Material and Methods

### Molecular cloning of FlyPer constructs

The genetic sequences for cyto-HyPer7, NLS-HyPer7 and mito-HyPer7 were amplified by high-fidelity PCR from plasmids generated in a previous study^11^. Briefly, to target HyPer7 to the cytosol, the nuclear export signal from the HIV-1 Rev protein was added to the C-terminus of HyPer7. For nuclear localisation, a triple nuclear localisation signal of simian virus 40 large T antigen was incorporated at the C-terminus of HyPer7. To target HyPer7 to the mitochondrial matrix, two COX8 pre-sequences were inserted at the N-terminus of HyPer7. Using In-Fusion cloning (Takara Bio), HyPer7 sequences were cloned into pCopia vectors (at NotI and BamHI restriction sites) for expression in cell culture, and into a pUAST-attB expression vector (at HpaI restriction sites) for the generation of transgenic FlyPer lines. The Myr-HyPer7 sequence^88^ was cloned into the pUAST expression vector^38^ (at EcoRI and XhoI restriction sites) for the generation of transgenic PM-FlyPer lines. All plasmid sequences were fully verified by Sanger sequencing.

### Generation of FlyPer lines

Embryo injection and generation of FlyPer lines were performed by BestGene, Inc (Chino Hills, CA, USA). Lines were generated using phiC31-mediated integration at an attP docking site on the third chromosome (BDSC #8622). The UAS-myr-HyPer7 construct was randomly inserted, and for this study a line with integration on the third chromosome was used.

### Fly strains and husbandry

Experimental flies were kept at 25°C on a 12 h light:dark cycle with constant humidity. Standard fly medium typically consisted of sugar-yeast-agar (SYA) diet consisting of 5% w/v sucrose (Sigma-Aldrich), 10% w/v brewer’s yeast (MP Biomedicals, #903312), 1.5% w/v agar (Sigma-Aldrich), and with 30 mL/L of 10% w/v nipagin (Sigma-Aldrich) in 95% EtOH and 0.3% v/v propionic acid (Sigma-Aldrich) added as mould inhibitors when the food had reached a temperature below 60°C. For studying the larval wing discs and developing embryos, flies were reared on a Cornmeal diet (9% dextrose w/v, 7.5% maize w/v, 2.2% yeast w/v 1% agar w/v, and 30ml of nipagin per litre). In addition to the FlyPer lines generated in this study (see section “Generation of FlyPer lines”), the following strains were used: *white Dahomey* (*w^Dah^*) and daGAL4 (kindly donated by Helena Cochemé). Tubulin-GAL4 (#5138) and H2Av-RFP (#23651) were obtained from the Bloomington Drosophila Stock Center (NIH P40OD018537). FlyPer lines and driver strains used to study ageing, development time, viability and *in vivo* oxidation by paraquat were all backcrossed for at least 6-10 generations into the outbred *w^Dah^* wild-type strain.

### Cell culture

*Kc167* (*Kc*) cells were cultured in serum-free HyClone™CCM3 medium (Cytiva) at 27°C. To express HyPer7 constructs, *Kc* cells were plated at a density of 200,000 cells and were transiently transfected with 400 ng of purified plasmid using the TransIT-2020 transfection reagent (Mirus MIR5400) and according to the manufacturer’s instructions. After ∼72 h, cells were transferred in volumes of 100 µL to an 8-well µ-slide (IBIDI) and imaged at an LSM880 confocal microscope (Zeiss).

### Viability and development assay

Virgin daGAL4 females were crossed to UAS-HyPer7-NES, UAS-COX8-HyPer7, UAS-myr-HyPer7, UAS-HyPer7-3xNLS or *w^Dah^* males. Embryos from a synchronised period of egg laying (∼4 h) were gently transferred from apple juice plates to vials containing SYA food (n∼150-180 embryos per genotype, ∼30 embryos per vial). Development time until eclosion was monitored at regular intervals and the total number of enclosed adults was scored to calculate percentage viability for each genotype.

### Lifespan assay

To generate flies ubiquitously expressing cyto-HyPer7, female daGAL4 virgins were crossed to UAS-HyPer7-NES males or *w^Dah^* males for the control cohort. To synchronise development, embryos were collected on apple juice plates within an ∼8 h time window and transferred to SYA food at equal densities. The resulting offspring was allowed to mate for ∼48 h before females were selected and distributed over 10 vials in groups of 15. Every 2-3 days, flies were gently flipped, without anaesthesia, onto fresh food while the number of deceased flies per vial was recorded simultaneously.

### Paraquat feeding experiments

Methyl viologen dichloride hydrate (Paraquat; Sigma-Aldrich) was added to the standard SYA diet at a final concentration of 20 mM once the fly medium had cooled below 60°C. Female flies ubiquitously expressing mito-HyPer7 (daGAL4>UAS-COX8-HyPer7) were allowed to mate for ∼48 h, and were subsequently separated from males and aged for 7 days on regular SYA food. Flies were then flipped to vials containing either their standard diet or 20 mM paraquat-supplemented food. For 24 h, flies were allowed to feed before tissues were dissected (see “Tissue dissections”).

### Tissue dissections

Since all dissections were followed by immediate live confocal imaging, no more than two flies were dissected at a time, with multiple batches processed sequentially within one experiment. This was a precaution to minimise the risk of tissues artificially oxidising due to potential prolonged exposure to *ex vivo* conditions. For dissections of adult gut, fat body tissue, and malpighian tubules, female flies were anaesthetised by briefly placing them in an empty vial on ice (∼10-30s). Flies were then decapitated and transferred to a silicone-bottomed dissection dish containing HyClone™CCM3 medium (Cytiva). The gut, with malpighian tubules still attached, was dissected by gently pulling it out from the posterior end of the female abdomen, taking care to keep the intestine intact along its entire length. The fat body was dissected by cutting the abdomen open along the ventral midline using microscissors. All internal organs were then carefully removed and the abdomen gently detached from the thorax. Fat body tissue was then imaged by confocal microscopy while still attached to the abdominal cuticle. Tissues were typically mounted in 5-10 µL of CCM3 medium on a ColorView^TM^ adhesive microscope slide (StatLab), covered by a #1.5 coverslip (Epredia) and sealed with a layer of Halocarbon oil 700 (Sigma-Aldrich).

Wing imaginal discs were dissected from third instar larvae and mounted in Shields and Sang M3 Insect Medium (Sigma-Aldrich S8398) into 35 mm glass-bottomed chambers (Ibidi). Embryos were collected on apple agar plates (apple juice with 1.5% agar) after laying for approximately 3h. Embryos were dechorionated in 50/50 bleach/water for 3 min. Embryos were washed with water and mounted in a coverslip and a custom-made slide holder and covered with halocarbon oil to avoid desiccation and imaged for 15-16h in 20 min intervals.

Egg chambers were dissected as previously described^89^, where in brief, FlyPer females were separated 2-3 days prior to imaging, and vials were supplemented with additional yeast paste. Females were anaesthetised with CO_2_ followed by 20s of 70% ethanol submersion. Ovarioles were dissected in Schneider’s Insect medium (BioWest), supplemented with 10% Heat-inactivated Foetal Bovine Serum (Cytiva) and 1% Penicillin-Streptomycin (Thermo Fisher Scientific) at room temperature, and egg chambers were carefully separated. Stage 9 and Stage 10 chambers were chosen and transferred into 35 mm glass-bottomed chambers (Ibidi) containing Schneider’s Insect Medium supplemented with 200 μg/mL human insulin (Sigma-Aldrich I9278) . Chambers were imaged immediately after for 30 min at 0, 6 and 30 min time points.

### Tissue staining

To stain nuclei in the adult midgut, the tissue was incubated with 1:1000 Hoechst nucleic acid stain (Invitrogen) for ∼5 min at room temperature immediately after dissection and prior to live confocal imaging. For labelling of the plasma membrane, a freshly dissected adult midgut was incubated in 5 µg/mL FM^TM^4-64 dye (Thermo Fisher Scientific) in PBS for 1-2 min on ice and then imaged directly afterwards in the staining solution using live confocal imaging. To visualise mitochondria, we incubated adult malpighian tubules, while still attached to the gut, with 500 nM of MitoTracker Deep Red FM (Thermo Fisher Scientific) in PBS for ∼1 h at room temperature and protected from light. The tissue was then rinsed in PBS twice, before being captured by live confocal imaging.

### Ex vivo oxidation of tissues

*Ex vivo* oxidation of gut tissue was carried out by incubating the dissected tissue for 15 minutes in 250 µM H_2_O_2_ in PBS or 1 mM diamide in PBS, while protected from light. Tissues were then washed once in PBS, mounted on a slide in CCM3 medium, followed by live confocal imaging. *Ex vivo* oxidation of egg chambers was carried out by addition of 10 or 100 µM H_2_O_2_ in Schneider’s Insect Media whilst being imaged.

### Confocal imaging of FlyPer lines

All microscope settings were kept identical between conditions and age groups within each experiment. In addition, different conditions (e.g. paraquat vs control) and age groups (young vs old) were typically imaged within the same session on the same day.

Images of larval wing discs and egg chambers were taken using an Olympus spinning disk confocal microscope using the SoRa disk with Air UPLXAPO 20x 0.8 and oil UPLXAPO 40X1.4 objectives. Egg chambers, wing discs and PM-FlyPer and NLS-FlyPer embryos were imaged and excited sequentially using low laser power (1.5% - 4%), an exposure time of 100 ms and with a 0.35 µm Z interval (embryos and wing discs) or 3-4 µm interval (egg chambers). Mito-FlyPer and cyto-FlyPer embryos were imaged using a Nikon N1-AR confocal microscope and a 20x/0.75 Plan Apo objective, using 0.5% laser power and 0.35µm Z interval.

Confocal imaging of the adult gut and fat body was performed on an LSM880 confocal microscope (Zeiss) using ZEN 2.3 Sp1 (version 14.0.20.201) software. Samples were typically imaged using a 40x oil immersion objective (gut) or a 20x dry objective (fat body) and excited sequentially by 405 nm and 488 nm lasers at low laser power (∼0.2%). Z-stacks were imaged with a 0.5 - 1 µm interval. Since the *Drosophila* midgut is highly compartmentalised^90^, the anterior midgut (R1/R2 area) was imaged to maintain consistency between conditions and age groups.

### Image analysis

Image analysis was performed using FIJI (v2.16.0/1.54p^91^). Ratiometric images were generated according to a previously described protocol^80^. Briefly, for confocal images acquired on the LSM880, both channels were thresholded in FIJI using the Triangle algorithm, with manual adjustment for each image, and background pixels set to NaN to exclude pixels lacking sufficient signal in either channel from the ratio calculation. The (thresholded) 488 nm image was then divided by the thresholded 405 nm image using the image calculator tool to generate a 32-bit floating-point result. Ratiometric images were then colour coded with either the Fire or mpl-Magma LUT. For images acquired using the spinning disk microscope, the 488 nm channel was divided directly by the 405 nm channel without thresholding.

For quantitative mito-FlyPer and cyto-FlyPer analysis of the adult midgut, regions of interest (ROI) were manually drawn around individual enterocytes (∼20-30 cells per gut) on a single Z-slice, typically selecting the optical section closest to the cell midplane. Since cyto-HyPer7 relies on active nuclear export, an additional ROI was drawn around the nucleus to exclude residual nuclear signal from measurements. HyPer7 oxidation ratios were measured for each cell by dividing the background-corrected 488 nm excitation intensity (oxidised) by the 405 nm excitation intensity (reduced). As individual cells were more difficult to distinguish in the fat body, two locations were imaged per tissue and the mean oxidation value per Z-slice was obtained. Subsequently, the mean oxidation value per Z-stack was calculated and the two means were averaged to obtain one oxidation value per tissue. For quantitative mito-FlyPer analysis of egg chambers, ROIs were manually drawn around the whole chamber on the max-projection ratio image stack and the raw integrated density was divided by the area of the ROI to generate the oxidation value at each time point.

Statistical analyses were performed in GraphPad Prism (version 11.0.2). ImageJ and Adobe Photoshop were used to rotate and resize images, and to fill empty canvas with background.

## Supplemental information

Document S1. Figures S1–S3

Movies S1-S4. Embryogenesis timecourses.

