## Supplementary figures for "Mapping subcellular H_2_O_2_ dynamics reveals tissue specific redox patterns in *Drosophila*"

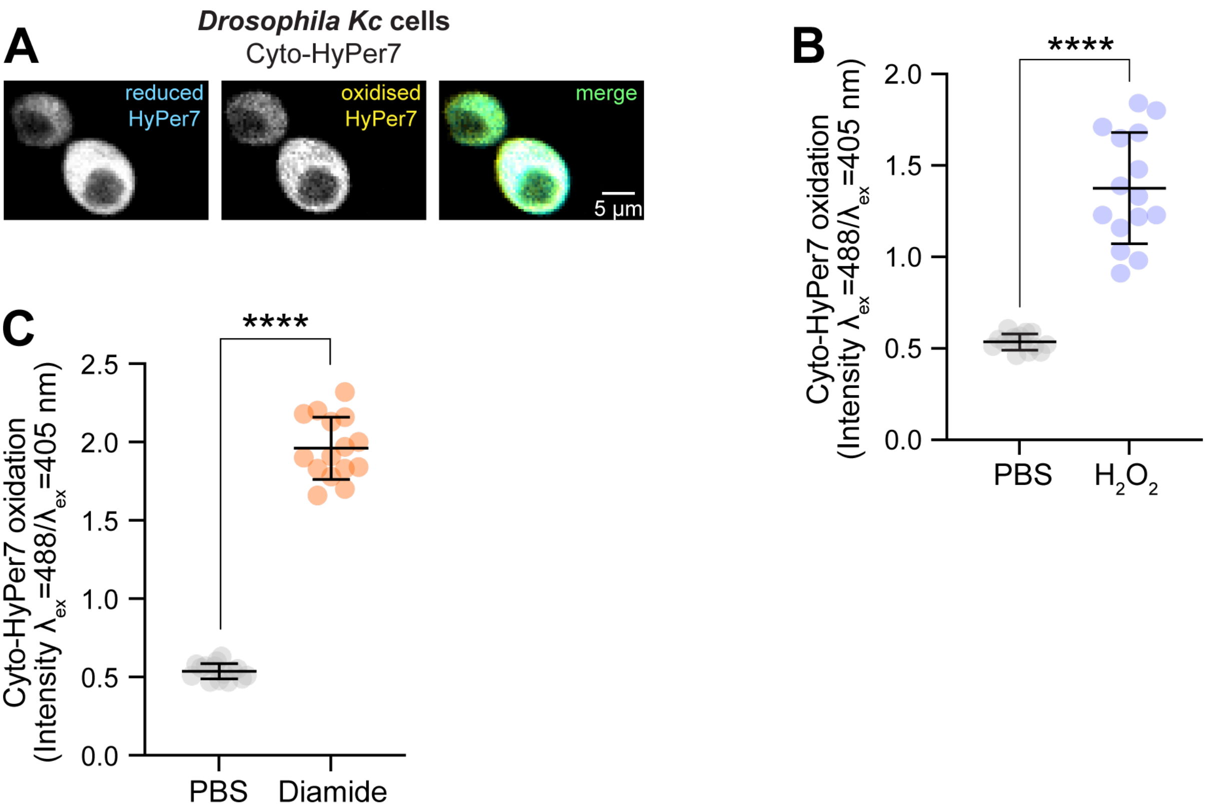


**Figure S1. HyPer7 can be expressed in *Drosophila Kc* cells and is functional.**

**(A)** Representative confocal image of *Kc* cells expressing cytosolic HyPer7 (cyto-HyPer7). Images show reduced HyPer7 (left column), oxidised HyPer7 (middle column) and the merged signal (right column). Scale bar = 5 µm. **(B)** Quantification of cyto-HyPer7 upon exogenous $H_2O_2$ application (250 µM in PBS, 15 min treatment) compared to the control treatment (PBS, 15 min treatment). Data were analysed by Welch's t-test (****, $p<0.0001$). $n$ = 15 cells. **(C)** Quantification of cyto-HyPer7 upon exogenous diamide application (1 mM in PBS, 15 min treatment) compared to the control treatment (PBS, 15 min treatment). Data were analysed by Welch's t-test (****, $p<0.0001$). $n$ = 15 cells.

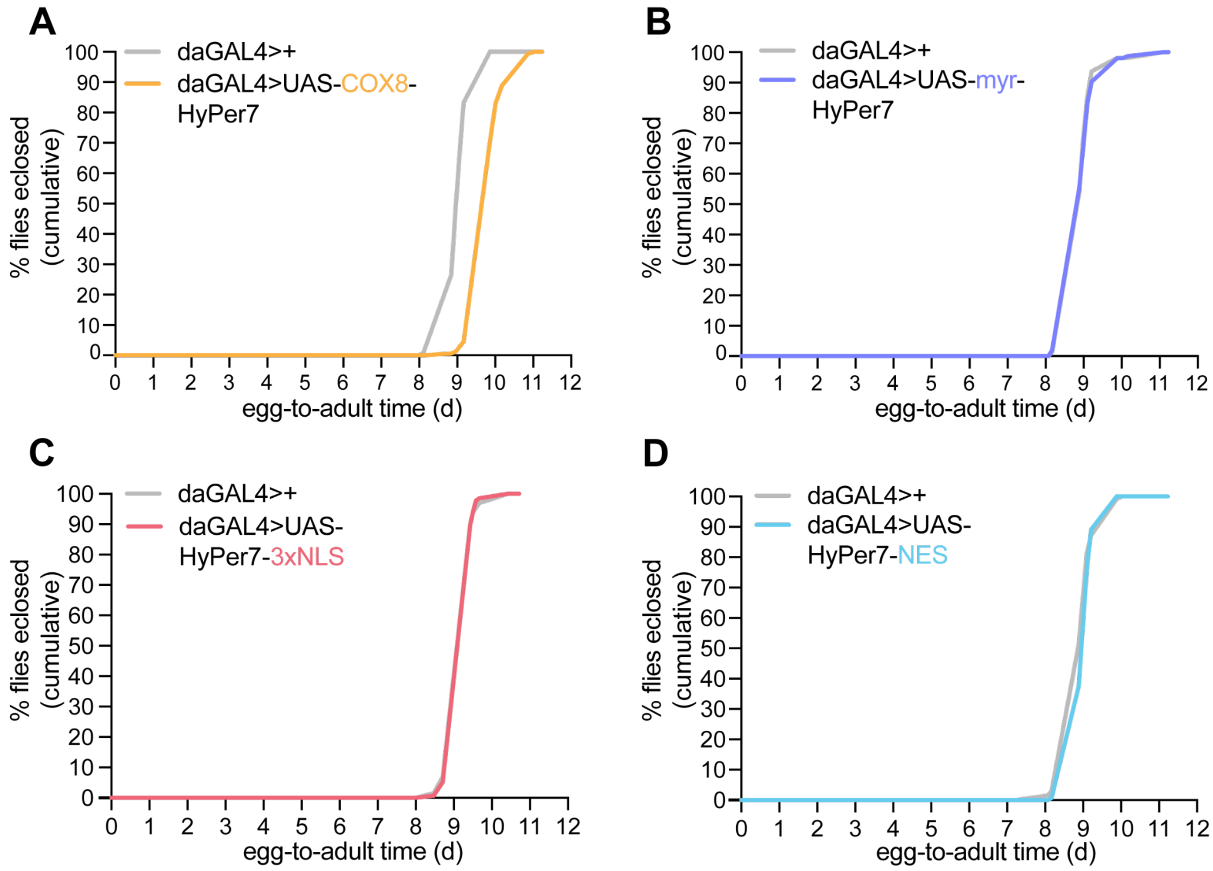


**Figure S2. Development from egg-to-adult of FlyPer lines ubiquitously expressing HyPer7.**

**(A-D)** Development time from embryo to eclosion of the different FlyPer lines ubiquitously expressing mito-HyPer7 (A), PM-HyPer7 (B), NLS-HyPer7 (C) or cyto-HyPer7 (D) compared to controls (daGAL4/+). Data are presented as the mean cumulative eclosion percentage. *n* = 5–6 vials per genotype, each containing 25-30 embryos.

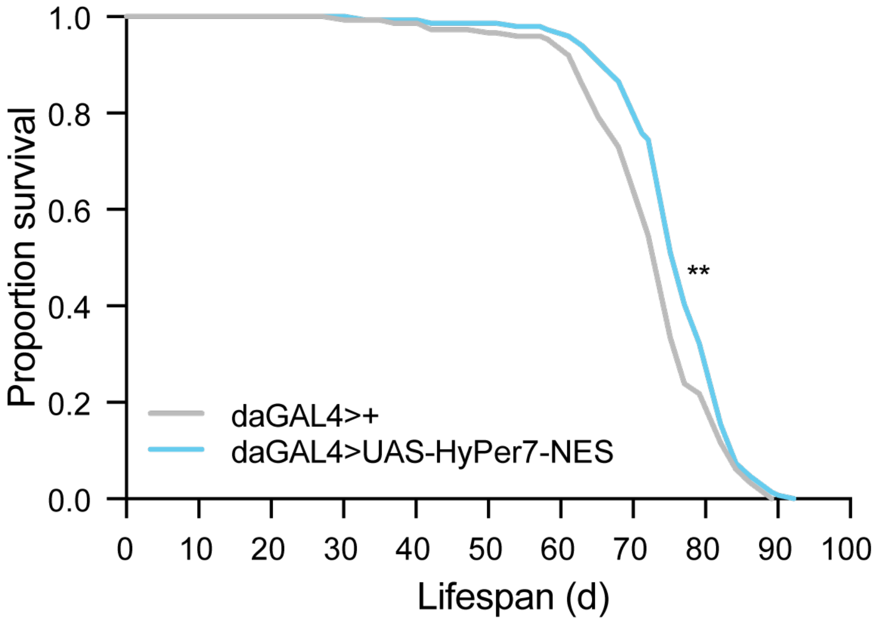


**Figure S3. Lifespan of female flies ubiquitously expressing cytosolic HyPer7 compared to controls**

Lifespan curve of flies ubiquitously expressing cytosolic HyPer7 (daGAL4>UAS-HyPer7-NES) and control flies (daGAL4>+). Statistical analysis was performed using Log-Rank test (**, $p < 0.01$). $n = 150$ female flies per condition.
